# A genotype-phenotype transformer to assess and explain polygenic risk

**DOI:** 10.1101/2024.10.23.619940

**Authors:** Ingoo Lee, Zachary S. Wallace, Yuqi Wang, Sungjoon Park, Hojung Nam, Amit R. Majithia, Trey Ideker

## Abstract

Genome-wide association studies have linked millions of genetic variants to biomedical phenotypes, but their utility has been limited by lack of mechanistic understanding and widespread epistatic interactions. Recently, Transformer models have emerged as powerful machine learning architectures with potential to address these and other challenges. Here we introduce the Genotype-to-Phenotype Transformer (G2PT), a framework for modeling hierarchical information flow among variants, genes, multigenic systems, and phenotypes. As proof-of-concept, we train G2PT to model the genetics of metabolic traits including insulin resistance (serum triglycerides-to-HDL ratio), LDL and type-2 diabetes. G2PT predicts these traits with accuracy exceeding state-of-the-art and, unlike other polygenic models, extends to distinct populations not used for training. Predictions of insulin resistance are based on >1,395 variants within 20 systems and include epistatic interactions among variants, e.g. between *APOA4* and *CETP* in phospholipid transfer. This work positions hierarchical graph transformers as a next-generation approach to polygenic risk.

## Introduction

Common diseases such as type 2 diabetes (T2D)^1^, cardiovascular disease^2^, and fatty liver^3^ are highly polygenic and physiologically heterogeneous, involving complex networks of interactions within and among multigenic molecular systems^4^. In these complex traits, individual examination of single nucleotide polymorphisms (SNPs) and other genetic variants has had limited utility for risk prediction and stratification in most individuals. Rather, progress has been made by systematically scanning for associated SNPs through genome-wide association studies (GWAS)^5^, and then combining these many SNP-phenotype associations using methodologies collectively termed Polygenic Risk Scores (PRS)^6–8^. PRS methods have been applied to predict risk for a wide range of common multigenic diseases^1,9–11^ but have two major limitations. First, they model loci additively and thus miss functional dependencies among variants, including genetic epistasis. Second, they assign a summary risk score for the whole collection of SNPs comprising an individual’s genotype, without explaining how that risk is attributed to perturbations to molecular and physiological pathways, or recommending what action the individual should take. Addressing these aspects could markedly improve risk prediction, biological interpretation and ability to guide treatment^12^.

Recent developments in deep learning^13–15^ offer significant opportunities to advance the PRS framework^16–19^, as they have the capacity to model both complex epistatic interactions and knowledge of molecular mechanisms. Among these, the Transformer has shown strong potential to address longstanding problems in the biomedical sciences, including prediction of 3D protein structures^20–22^, biomedical image analysis^23^, inference of gene expression from genome sequence^24–26^, and mapping a sequence of SNPs to a predicted phenotype^27,28^. The Transformer architecture is known for its central use of an “attention” mechanism^29^, an operation that dynamically computes the importance of each input element relative to others, enabling the model to focus on the most relevant features^29–31^. While this mechanism can aid in interpretation, the vast number of input features typical of biomedical data can greatly complicate interpretability and transparency. To address this challenge, incorporating biological knowledge into Transformer attention – such as that provided by gene function databases like the Gene Ontology^32^ – has the potential to improve model explainability, transparency, and fairness, all of which are crucial for clinical applications^33–38^.

Here we describe the Genotype-to-Phenotype Transformer (G2PT), a graph transformer architecture for general genotype-to-phenotype translation and interpretation (**Fig. 1a**). The G2PT model analyzes the complex set of genetic variants in a genotype by directing attention to their effect on embedded representations of genes and a hierarchy of multigenic systems. As proof-of-concept we use G2PT to study metabolic traits including insulin resistance (as measured by the triglyceride to high-density lipoprotein cholesterol ratio: TG/HDL), serum cholesterol (low-density lipoprotein cholesterol: LDL) and T2D^39,40^, the most common metabolic disease. Hundreds of loci have been mapped for each of these (TG/HDL^41,42^, LDL^43^, T2D^44,45^) but the genetic circuits that regulate their status and molecular physiology are still poorly understood. In what follows, application of G2PT yields a predictive genomic model that explains metabolic traits via a constellation of genetic factors, biological systems, and nonlinear epistatic interactions.

**Fig 1:**
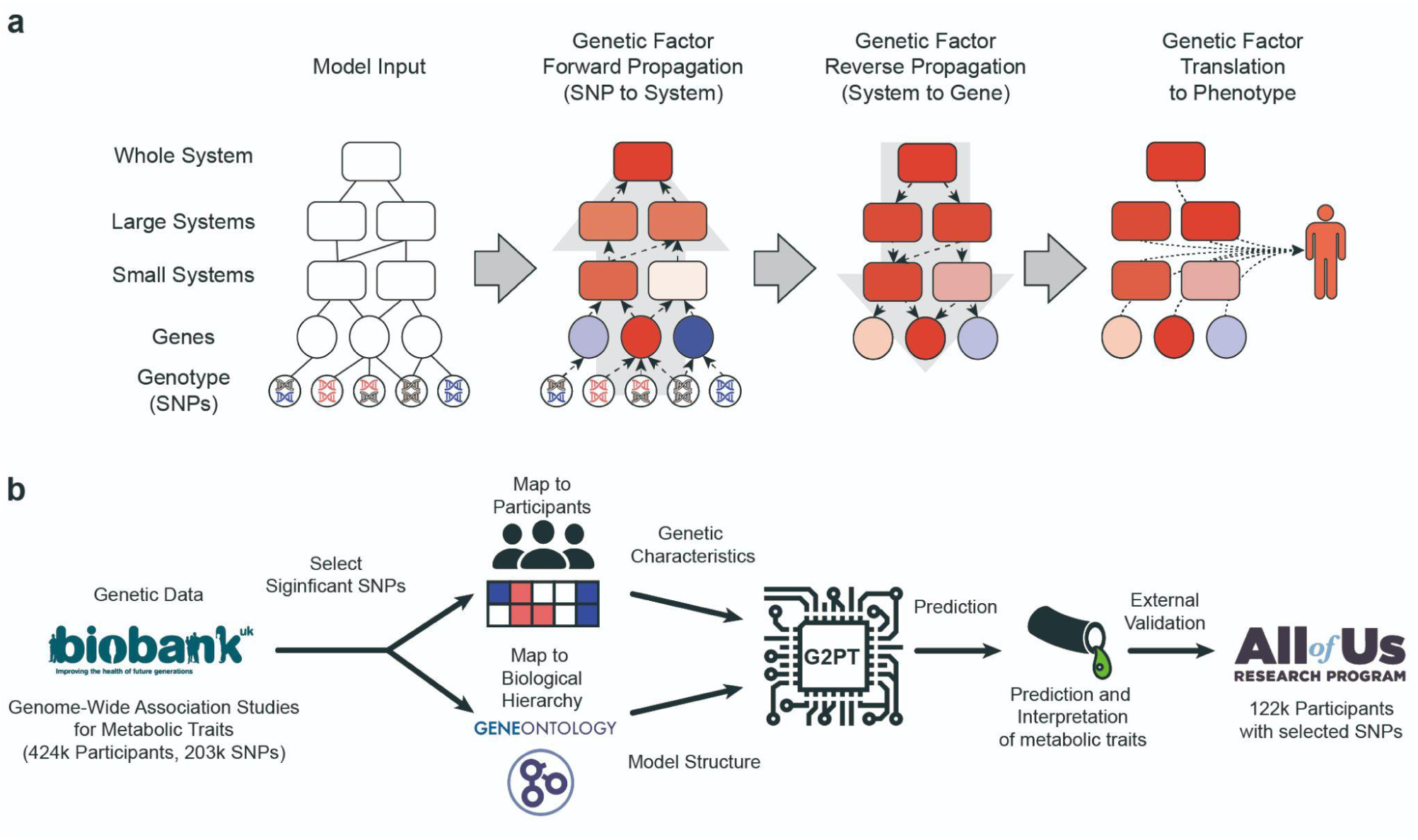
G2PT workflow. **a,** Inputs of the Transformer model include genotypic data (SNPs, bottom layer), a mapping of SNPs to genes (second layer), and a mapping of genes into a hierarchy of multi-genic molecular systems (top layers). The presence of SNP minor alleles modifies the embedding states of downstream genes and multigenic systems (forward propagation). Conversely, state changes in systems influence the states of sub-systems and genes they contain (reverse propagation). Finally, all gene and system states are integrated to predict phenotype. **b,** Proof of concept via prediction of metabolic traits such as triglyceride/HDL cholesterol ratio (TG/HDL), low-density lipoprotein cholesterol (LDL), and type-2 diabetes (T2D). Genotypic data and corresponding metabolic traits are extracted for participants from the UK Biobank. SNPs are selected based on their independent association with metabolic traits and mapped to the closest genes, which in turn map to multigenic systems defined by the Gene Ontology. This information is used by G2PT to predict metabolic phenotypes.

## Results

### G2PT Model Overview

The G2PT framework models the states of biological entities, including variants, genes, multigenic systems, and phenotypes, as coordinates within a machine learning embedding. An embedding is a simplified low-dimensional representation of a high-dimensional dataset, optimized so that similar entities are assigned similar embedding coordinates^46^. Positions in the embedding (i.e. the states of each entity) are governed by a hierarchical graph transformer, a deep neural network that models flow of information across a hierarchy of connected entities. Such information flow includes the effects of variants on the states of genes (SNP-gene mapping), the effects of altered genes on multigenic systems and their supersystems (gene-system and system-system mapping), and the reciprocal influences of systems on the states of their component systems and genes (**Fig. 1a**, **Methods**). Based on the collection of variants comprising an individual’s genotype, the model uses a multi-head attention mechanism to propagate these effects to select biological entities in the hierarchy, resulting in updates to their embedding coordinates. Finally, the entire collection of embedding states for genes and systems is used to predict phenotype.

### Using G2PT to Model Metabolic Traits and Disease

As proof-of-concept, we used G2PT to study human metabolism, focusing on insulin resistance (TG/HDL), LDL cholesterol, and T2D (**Fig. 1b**). Serum TG, HDL, LDL, and T2D case:control status were obtained from 423,888 participants profiled in the UK Biobank with accompanying genetic data (covering 203,126 SNPs; **Methods**, **Supplementary Fig. 1**)^42,47^. SNPs were mapped to genes using any of three lines of evidence: expression quantitative trait loci (eQTL)^48^, curated mappings from the cS2G mapping tool^49^, or the closest gene in genomic coordinates (hg37 reference genome, **Methods**). Genes were mapped to a hierarchy of multigenic systems condensed from Biological Process terms recorded in the Gene Ontology (**Fig. 1b**)^32^.

Using this information, G2PT models were trained to translate an individual’s complement of SNP alleles into a prediction of each of their three metabolic trait values (separate models for TG/HDL, LDL, T2D). Training and evaluation were carried out in a robust framework of five-fold nested cross-validation^46^ (**Methods**). Input features for G2PT were defined as SNPs that have an independent marginal association with metabolic phenotypes, where significance of association was defined across a series of p-value thresholds of decreasing stringency (models built on SNP sets ranging from p<10^−8^ to p<10^−4^, **Methods**, **Supplementary Table 1**). The entire training procedure across all models required approximately 10 hours using 4 NVIDIA A30 Graphics Processing Units (GPUs, **Methods**).

Following model training, we assessed the performance of G2PT in predicting the three metabolic traits for held-out UK Biobank individuals. Performance was benchmarked against a panel of widely used PRS and machine learning approaches. This panel included PRS methods based on SNP thresholding or thresholding and clumping (PRS T, PRS C+T, PRScs with threshold, SBayesRC with threshold)^50^; regularized linear and nonlinear regression models (ElasticNet, XGBoost)^51,52^; and polygenic scoring methods based on the entire genome–wide collection of SNPs (Lassosum, LDpred2, PRScs, SBayesRC^53–56^). Across all SNP p-value thresholds, G2PT achieved an explained variance (R^2^) that was significantly higher than either PRS or regularized regression approaches (**Fig. 2a–c**). Notably, this performance advantage persisted even in comparison to advanced models utilizing genome-wide summary statistics (LDPRED2, Lassosum2, PRScs, and SBayesRC). G2PT outperformed these dense models despite relying on only thousands of prioritized SNPs, in contrast to the hundreds of thousands of genome-wide approaches (**Fig. 2a–c**, right panels).

**Figure 2.**
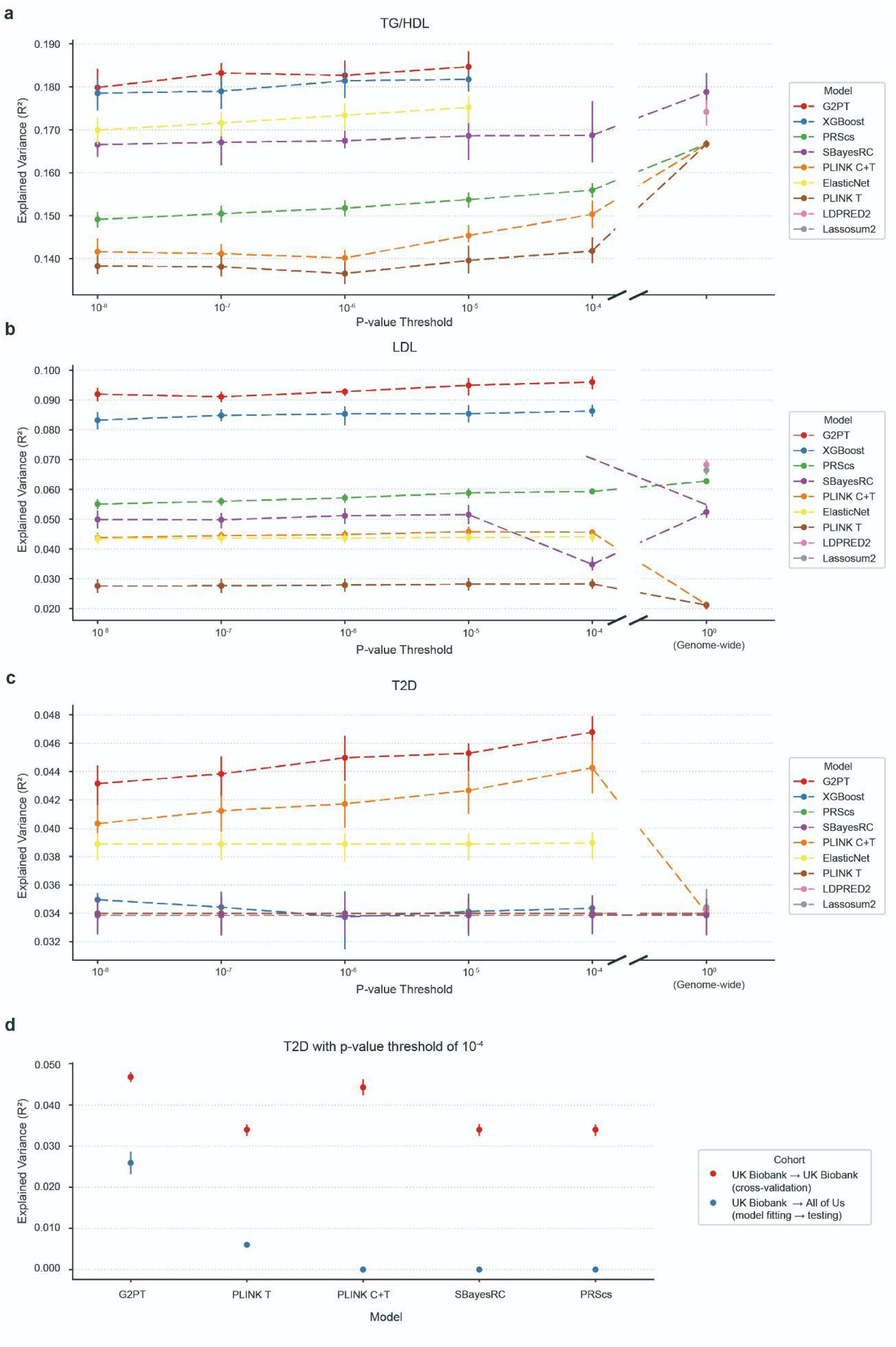
Predictive performance of G2PT compared with PRS models. **a**, Explained variance (R^2^) of prediction of TG/HDL phenotype. The left section of each panel compares G2PT against ElasticNet, XGBoost, PRS clumping + thresholding (PLINK C+T), PRS thresholding (PLINK T), PRScs, and SBayesRC, across varying marginal p-value thresholds for SNP selection and corresponding number of SNPs. The right section (after the axis break) compares performance against various genome-wide PRS methods: PLINK T, PLINK C+T, LDPRED2, Lassosum2, PRScs, and SBayesRC. **b–c**, As for panel (**a**) but for LDL and T2D phenotypes, respectively. **d**, Validation of T2D prediction performance (R^2^) using a p-value threshold of 10^−4^. Models are trained on the UK Biobank and performance is evaluated either in a held-out partition of UK Biobank (red) or on the All of Us cohort (blue) to assess transferability. Points represent mean performance over nested five folds of cross-validation, with error bars showing 95% confidence intervals.

In addition to cross-validation studies within the UK Biobank, we also evaluated the degree to which predictions of the G2PT model were transferrable to other human populations not seen during training. For this purpose we obtained genotype-phenotype data from the All of Us cohort^57^, comprising 122,340 individuals of whom 41,849 had a prior diagnosis of T2D (covering 706 SNPs, **Fig. 1b**). When trained in the UK Biobank and evaluated on this All of Us cohort, G2PT retained significant predictive power (R^2^=0.025) albeit lower than seen in cross-validation (**Fig. 2d**). Notably, all competing methods tested suffered severe performance degradation, losing most to all predictive power when transferred to All of Us for T2D case:control prediction (**Fig. 2d**).

To investigate which aspects of the G2PT model were most responsible for the increased performance, we repeated our analysis over a series of ablation studies in which key G2PT architectural modules were removed. These studies included [1] removing knowledge of multi-gene systems, thereby training the Transformer with SNP inputs and SNP-to-gene mappings only; [2] removing knowledge of both systems and genes, leaving only SNPs; and [3] removing hierarchical information flow between systems and supersystems, treating each system as an independent entity. The full G2PT model outperformed all of these ablations either modestly (most comparisons) or substantially (removal of systems and genes), indicating that incorporating hierarchical biological knowledge does not limit predictive capability compared to simpler black-box approaches (**Supplementary Fig. 2**). We also examined the effects of simplifying our SNP-to-gene mapping policy. We found that removing the more advanced eQTL and cS2G mapping methods, leaving a model based on the closest gene only, led to a significant albeit modest decrease in prediction performance (**Supplementary Fig. 2**). In addition, we quantified performance as a function of training set size (number of individuals) by subsampling the cohort and retraining G2PT with identical architecture and SNPs (**Supplementary Fig. 3**). We found that stable performance emerges by ∼100k samples for the continuous traits (TG/HDL, LDL) but continues to improve for the binary trait (T2D), consistent with lower per-sample information in case–control settings.

### Transformer Attention Reveals Genes and Mechanisms Underlying Phenotype

We next turned to mechanistic interpretation – studying the model’s transformer attention mechanism to reveal the key genes and systems it had used for prediction, focusing on TG/HDL (**Methods**). As a positive control, we examined the lipoprotein lipase (*LPL*) gene, a known insulin resistance gene ^58^ which we expected should be given high attention by the G2PT framework. We indeed found participants for which G2PT gave high attention to variants in *LPL*, and that these individuals tended to have high predicted TG/HDL (**Supplementary Fig. 4a**). A similar relationship was observed for the molecular systems in which this gene is involved (e.g., lipid localization, **Supplementary Fig. 4b**).

Across all TG/HDL predictions for the 420K+ individuals, we identified significant model attention on 20 multigenic systems incorporating a total of 1,395 SNPs linked to 253 genes (**Fig. 3a**, **Methods**). Of these genes, 172 had been associated with TG/HDL ratio in our most recent GWAS^42^, whereas the remaining 81 had not (**Fig. 3a**). In general, model attention on systems was higher than on genes (**Supplementary Fig. 4c**) and not significantly correlated with system size (number of genes), depth in the systems hierarchy, or gene-level importance (**Supplementary Fig. 4d-f**). Key systems included cholesterol and phospholipid transport, lipid storage, vesicle docking, and transcriptional regulation – corresponding to known metabolic pathways for serum lipids and their accompanying regulatory factors (**Fig. 3a**, **Extended Data 1**). We compared these results with MAGMA gene set analysis^59^, finding moderate agreement between the Gene Ontology systems identified by the two approaches (ρ = 0.26, p < 0.01**, Supplementary Table 2**).

**Fig 3:**
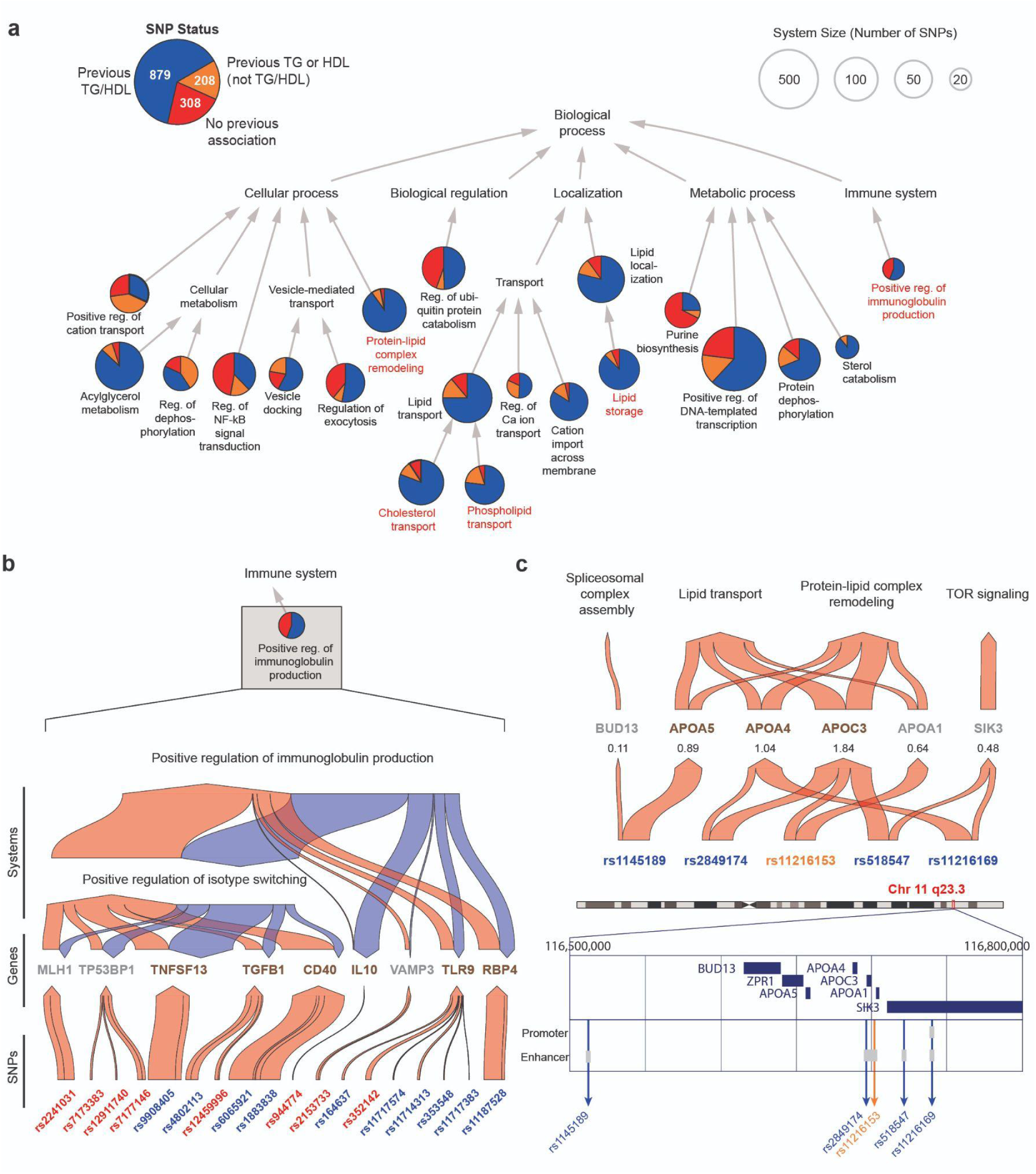
Multigenic systems determining prediction of TG/HDL. **a**, Hierarchy of important multigenic systems. Top 20 systems are represented as circular nodes, with size proportional to the number of genes annotated to the system. Arrows represent involvement in broader systems (Gene Ontology “is_a” relation) or containment of one system by another (“part_of” relation). Pie charts show the proportion of genetic variants assigned to each system in three categories: SNPs previously associated with TG/HDL ratio (blue); SNPs associated with TG or HDL individually (orange); SNPs not previously reported for any of these phenotypes (red). The pie chart in the legend (upper left) shows the total number of variant counts across the systems hierarchy. Red labels denote systems highlighted in subsequent figures. **b**, Genetic information flow within Positive regulation of immunoglobulin production (GO:0002639), a top-20 informative system. Red arrows represent information flow from SNPs to genes to systems (ascending layers). Blue arrows indicate flow from systems back to genes (reverse propagation). Arrow width is proportional to the amount of attention given by the model. SNPs previously associated with TG/HDL ratio are highlighted in blue; SNPs not previously associated with TG, HDL, or TG/HDL phenotypes are highlighted in red. Genes implicated due to a ‘closest gene’ mapping policy are colored in gray, whereas genes implicated due to other SNP-to-gene mappings (**Methods**) are dark brown. **c**, Genetic information flow at chromosomal locus 11q23.3. Top: genetic information flow visualized similarly to panel b, focusing on forward propagation of information only. SNPs previously associated with TG/HDL ratio are highlighted in blue; SNP previously associated with HDL is highlighted in orange. Shown beneath each gene is the sum of attention scores from its potential SNP-to-gene mappings. Bottom: Positions of SNPs with respect to the chromosomal locus. The whole chromosome 11 ideogram is shown, with coordinates 116.5 – 166.8 MB expanded to detail gene open reading frames (dark blue rectangles) and regulatory regions (promoters or enhancers; light gray).

G2PT highlighted several systems not emphasized in previous TG/HDL gene mapping studies or in the MAGMA results, suggesting hypotheses for further investigation. In particular, immunoglobulin production, the fundamental process of adaptive immunity which involves the creation of antibodies by B cells in response to antigens, was broadly supported by model attention to 18 SNPs with effects distributed over 9 genes (**Fig. 3b**). Supporting this observation, clinical investigations of individuals with immunoglobulin-related deficiencies report alterations in serum lipids and insulin resistance^60^. Further investigation revealed that ten of the SNPs impacting this system had in fact been associated with TG/HDL previously^42^, but independently from one another without any indication of a common biological process.

For SNPs with several potential SNP-to-gene mappings (**Methods**), we found that G2PT often prioritized one of these genes in particular due to its membership in a high-attention system. For example, the chr11q23.3 locus contains multiple genes including the *APOA1/C3/A4/A5* gene cluster^61^ (**Fig. 3c**) which is well-known to govern lipid transport, an important system for G2PT predictions (**Fig. 3a**). Due to high linkage disequilibrium in the region, all of its associated SNPs had multiple alternative gene mappings available. For example, SNP rs1145189 mapped not only to *APOA5* but to the more proximal *BUD13*, a gene functioning in spliceosomal assembly^62^ (a system receiving substantially lower G2PT attention). Here, the relevant information flow learned by G2PT was from rs1145189 to *APOA5* to lipid transport and protein-lipid complex remodeling (**Fig. 3c**; and conversely, deprioritizing *BUD13* as an effector gene for TG/HDL). We found that this particular genetic flow was corroborated by exome sequencing^63,64^, which implicates *APOA5* but not *BUD13* in regulation of TG/HDL, using data that were not available to G2PT. Similarly, two other SNPs at this locus – rs518547 and rs11216169 – had potential mappings to their closest gene *SIK3*, where they reside within an intron, but also to regulatory elements for the more distant lipid transport genes *APOC3* and *APOA4.* Here, G2PT preferentially weighted the mappings to *APOC3* and *APOA4* rather than to *SIK3* (**Fig. 3c**). Altogether, of the 1,395 SNPs important to G2PT predictions, 252 were involved in multiple potential SNP-to-gene mappings where G2PT had prioritized one of these genes substantially above the others (>2-fold difference in attention). These findings demonstrate how modeling genetic information flow through a hierarchy of SNPs, genes and multigenic systems provides a means to identify the major molecular pathways underlying a phenotype.

### Model Predictions Invoke Epistatic Relationships

The favorable performance of G2PT in comparison to linear models (**Fig. 2**) suggested that its predictions may leverage nonlinear (epistatic) interactions among SNPs. To investigate, we initially focused on SNPs impacting phospholipid transport, a system that was highly important to G2PT predictions of TG/HDL (**Fig. 3a**) predominantly through the phospholipid efflux subsystem. SNPs in this system were first selected based on high model attention (**Fig. 4a**), from which pairs of SNPs spanning distinct loci were tested for pairwise interactions using a standard statistical model of epistasis (**Methods**). This screen yielded six SNPs involved in seven epistatic interactions (**Fig. 4b**). For example, we identified an interaction between SNPs rs11216169 and rs7499892 located at distinct chromosomal loci encoding the genes apolipoprotein A-IV (*APOA4*, 11q23.3) and cholesteryl ester transfer protein (*CETP*, 16q13), respectively. For both SNPs, the minor alleles were linked to an increase in TG/HDL (**Fig. 4c**). However, the impact of rs11216169 was significantly amplified in the rs7499892 T/T homozygous minor genotype, an effect which was not only seen in the observed TG/HDL ratio (**Fig. 5c**) but captured in the model prediction (**Fig. 4d**). Beyond this particular interaction, the *CETP* locus harbored a total of five SNPs exhibiting epistasis with *APOA4* (**Fig. 4b, Supplementary Table 2**). Notably, the *APOA4* and *CETP* gene products are secreted into blood (from intestine and liver respectively), in which they associate with HDL particles in regulation of cholesterol ester transport^65,66^, supporting a functional biochemical interaction.

**Fig 4:**
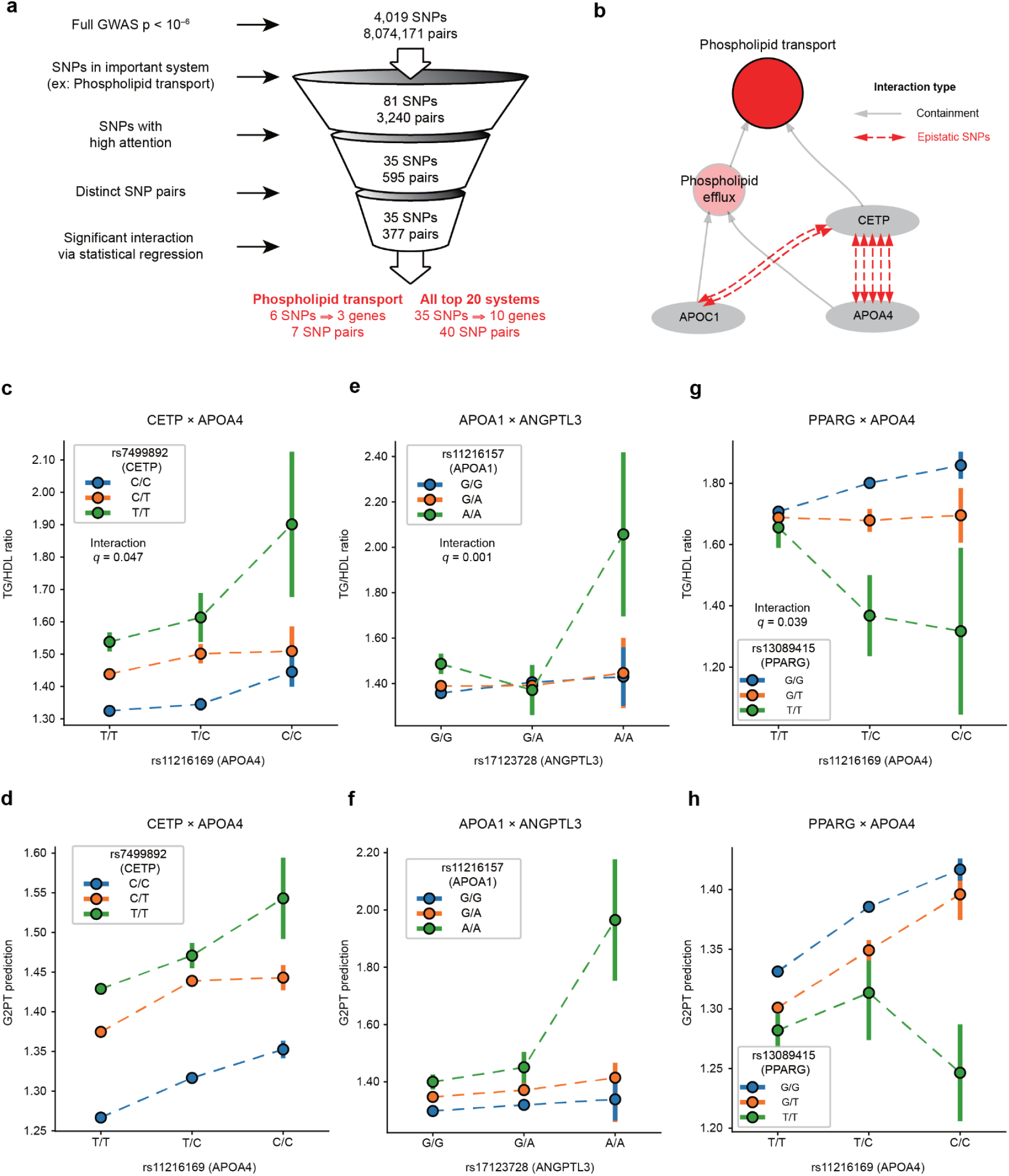
Epistatic interactions identified by attention-based epistasis search. **a**, Pipeline for detecting epistasis. Interacting SNP pairs are identified through a progressive multi-step screening process based on the system under study (here, Phospholipid transport), model attention and validation by statistical regression. **b**, Hierarchy of subsystems and individual gene products within phospholipid transport (red circle), with gray arrows representing containment of a gene product (gray ovals) or subsystem (pink circles) within a larger system. Red dashed arrows indicate significant epistatic interactions. **c**, Epistatic interaction of rs7499892 (CETP) with rs11216169 (APOA4) on measured TG/HDL ratio. Points and error bars show medians with standard error of TG/HDL ratio for subsets of individuals stratified by genotype. Colors denote rs7499892 genotype: C/C (blue, homozygous major allele), C/T (orange, heterozygous), and T/T (green, homozygous minor allele). X-axis denotes rs11216169 genotype: T/T (homozygous major allele), T/C (heterozygous), C/C (homozygous minor allele). **d,** Similar to panel (c), but showing the TG/HDL values predicted by the G2PT model. **e,f,** Epistatic interaction of rs17123728 (ANGPTL3) with rs11216157 (APOA1) on (e) measured or (f) predicted TG/HDL ratio. Display as per panel (c). **g,h**, Epistatic interaction of rs13089415 (PPARG) with rs11216169 (APOA4) on (g) measured or (h) predicted TG/HDL ratio. Display as per panel (c).

**Figure 5:**
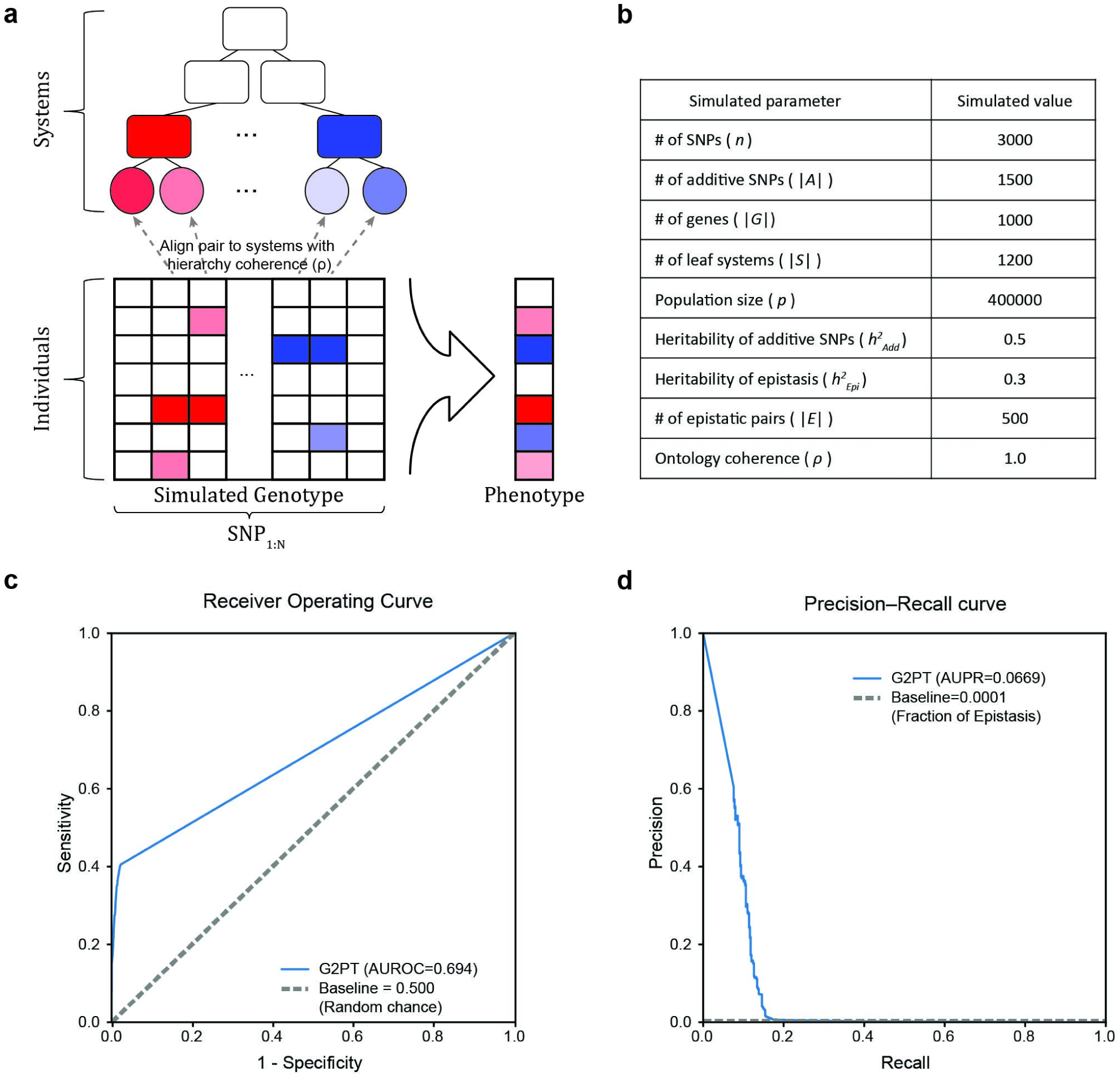
Simulations to explore detection of SNP-SNP epistasis. **a**, Schematic of the simulation. Standardized genotypes and phenotypes are initialized with specified numbers of additive SNPs and epistatic SNP pairs. A random system hierarchy is constructed, with epistatic SNPs assigned to systems based on an ontology coherence parameter ρ. Increasing ρ increases the likelihood that epistatic SNP pairs co-locate within the same leaf system. **b**, Table of parameters for epistatic simulation, selected for consistency with population studies in modern biobanks. **c**,**d**, Detection of epistatic SNPs is shown via ROC (**c**) and PR (**d**) curves. The random baseline is given by the fraction of true epistatic pairs.

Beyond phospholipid transport, we broadly applied this same epistasis screening procedure across all 20 systems most important to TG/HDL predictions (**Fig. 4a**). This broader screen identified 40 SNP-SNP pairs with significant epistasis (**Supplementary Table 3**). One significant interaction was identified in the lipid storage system, between variants at the apolipoprotein A-I (*APOA1*, 11q23.3) and angiopoietin-like 3 (*ANGPTL3*, 1qp1.3) loci; here phenotypic impact was seen predominantly in the combination of homozygous minor states for both variants (**Fig. 4e,f**). This interaction was also supported by the AoU cohort (**Supplementary Fig. 5**). As another example, screening of the cholesterol transport system again highlighted the *APOA4* gene, in which the rs11216169 minor allele exposed a strong suppressive effect of the rs13089415 variant upstream of peroxisome proliferator-activated receptor gamma (*PPARG* at locus 3p25.2, **Fig. 4g,h**).

### Validation of Epistasis through Simulation Studies

Having identified epistatic interactions in the study of metabolic phenotypes, we sought to characterize more generally G2PT’s ability to recover such non-additive effects under controlled conditions. For this purpose we developed a simulation–retrieval framework mirroring G2PT’s hierarchical architecture (**Fig. 5a**). In each simulation a random systems hierarchy was constructed, after which SNPs were assigned phenotypic effects of varying strengths as well as, for select SNP pairs, epistatic interactions. The choice of epistatic SNP pairs was governed by an ontology coherence parameter, ranging from ρ=1 (every epistatic interaction occurs between genes in a common leaf system) to ρ=0 (every epistatic interaction occurs between genes in two unrelated systems). Analysis of simulation results confirmed that epistasis detection accuracy increases consistently with ontology coherence (**Supplementary Fig. 6**). High ρ values improved recovery by concentrating epistasis within coherent systems, whereas low ρ values reduced epistasis detection to random chance. While absolute accuracy depended on other simulation parameters such as the sample size and number of epistatic interactions, G2PT maintained substantial improvement over random chance and over baseline methods such as PLINK^67^. At parameter settings consistent with current GWAS sample sizes and understanding of heritability^68,69^ (**Fig. 5b**), we observed a simulated AUROC of 0.694 (**Fig. 5c**), corresponding to recovery of approximately 10% of all epistatic relationships with >50% precision (**Fig. 5d**). Collectively, these simulations demonstrate that G2PT identifies epistatic interactions across diverse genetic architectures and clarify how detection strength scales with ontology coherence, sample number, and effect size.

## Discussion

Here we have introduced an approach to genotype-phenotype translation using a graph-based transformer (**Fig. 1**), a significant neural network architecture arising from recent machine learning research^70,71^. When applied to traits inducing insulin resistance (TG/HDL), serum LDL cholesterol, and T2D case status, this architecture outperforms genome-wide PRS methods (**Fig. 2**) despite being constrained to a few thousand loci. When trained in UK Biobank and evaluated in the ancestrally diverse All of Us cohort, G2PT retained measurable predictive power for T2D, whereas competing PRS-based methods exhibited near-complete performance collapse (**Fig. 2d**). This divergence is consistent with a well-recognized limitation of PRS, sensitivity to ancestry-specific linkage disequilibrium and allele frequency structure^72^, and it suggests that G2PT’s hierarchical aggregation of variant effects through genes and biological systems yields representations that are more robust to population heterogeneity.

Detailed examination of model attention identified a core set of biological systems driving phenotypic predictions, which integrate signals from hundreds of genes including expected and unexpected factors (**Fig. 3**). Among these was an unexpected system, immunoglobulin production, based on genotypic aggregation of 18 genetic variants impacting 9 genes (**Fig. 3a,b**). While some genes in this system were newly linked to metabolic phenotypes, others such as RBP4 are established biomarkers associated with insulin resistance in humans^73,74^ (but through distinct mechanisms from immunoglobulin production). Although few studies have directly linked insulin resistance to immunoglobulins, research in mouse models of diet-induced obesity suggests that B-cell infiltration into adipose tissue — producing pathogenic IgG — can drive inflammation and disrupt insulin signaling, ultimately contributing to systemic insulin resistance^75^. These observations suggest plausible mechanisms for further investigation in humans.

This ability of the G2PT model to identify key SNPs and systems was greatly aided by the formal mechanisms of attention inherent to Transformer architectures. During the ‘genetic factor propagation phase’ of modeling (**Fig. 1a**), we used multi-head attention to transmit the effects of SNPs across the knowledge hierarchy of genes and systems, allowing us to inspect and trace the cascade of molecular entities impacted by a genotype (**Fig. 3**). During subsequent ‘genetic factor translation’, single-head attention was used to quantify the impacts of the altered genes and systems on an individual’s phenotype. In this way, genetic factor propagation facilitates interpretation of the model by revealing complex information flows, whereas genetic factor translation integrates across these mechanisms to produce a single unified attention value used for interpreting the mechanisms underlying phenotypic risk.

A second notable aspect of the G2PT architecture is the bidirectional flow of information among genes and multigenic systems. Variant effects are first transmitted upwards in scale to impact genes and their collective functions, after which this flow is reversed to enable the functional states of systems to influence how specific variants impacting that system are interpreted (**Fig. 1a**). This reverse propagation step captures the biological context in which genes and variants operate (e.g., **Fig. 3b**), and it promotes cross-talk among multiple genetic variants that may have conditional interrelationships. These aspects enabled G2PT to learn gene-gene epistatic interactions across a variety of biochemical mechanisms. For example, in addition to interactions among apolipoproteins and their potential interactors (e.g. *APOA4* and *CETP*, **Fig. 4c**), G2PT identified novel epistasis between *ANGPTL3*, the major secreted inhibitor of lipoprotein lipase targeted by triglyceride-reducing therapies^76^, and *APOA1*, the major protein component of HDL particles (**Fig. 4e**). While these two gene products have not been reported to interact physically, recent data suggests they are transcriptionally regulated by the same hepatic nuclear factor *HNF1A*^77^, providing a potential mechanism for the observed genetic epistasis. Similarly, the identified epistatic interaction (**Fig. 4g**) between *APOA4*, an intestinally secreted triglyceride-rich lipoprotein factor, and *PPARG*, a nuclear hormone receptor and transcription factor, has a strong biological basis. The promoter of *APOA4* contains *PPAR* response elements, and *APOA4* expression is upregulated by the closely associated nuclear hormone receptor *PPARA*^78^. G2PT also found a non-canonical epistatic interaction between *TGFB1* and *FKBP1A* related to protein dephosphorylation (**Supplementary Table 3**). In particular, variants at the *TGFB1* locus can serve to increase TGF-β ligand–driven receptor/SMAD phosphorylation^79^, while variants at *FKBP1A* can weaken receptor activity^80^. Where both risk genotypes co-occur, this could plausibly lead to disproportionate SMAD activation, exaggerating transcriptional programs that raise TG and lower HDL, producing an epistatic increase in the TG/HDL ratio.

Despite the promise of Transformers for genotype–phenotype modeling, computational cost remains a principal bottleneck. Our use of the xFormer memory-efficient attention mechanism substantially reduced the memory footprint and accelerated training (**Supplementary Fig. 7**). Regardless, the current pipeline was capped at ∼5,000 SNP inputs before memory and interconnect overheads were found to dominate. We expect that parallelized, sharded execution can lift this limit in the near future, enabling larger SNP sets, faster convergence, and deeper hyperparameter exploration. We also experienced challenges arising from our use of Gene Ontology as a prior. This knowledgebase includes many redundant groups of systems with nearly identical sets of genes, and it has a natural bias towards well-studied systems^32^. An important step moving forward will be to explore alternative knowledge structures, such as Reactome^81^, GO Causal Activity Models (GO-CAMS)^82^ or maps of biological structures and systems resolved directly from ‘omics data^83–85^. Finally, our interpretability results relied on model attention, which we treat as hypothesis-generating rather than causal evidence. Correlations between attention and predictions do not establish mechanisms, and while we included stress tests (e.g., masking/ablation and permutation), such tests provide model-level, not biological, validation. Regardless, G2PT has immediate application to the genetic analysis of diverse phenotypes of interest, including those related to multigenic diseases such as autism, aging, or cancer. This work also has implications outside of the life sciences, insofar as it presents a general template for constructing interpretable Transformer architectures across deep learning challenges.

## Materials and Methods

### GWAS Data

Human genotype and phenotype data were obtained from the UK Biobank^47^, in which participants of Caucasian ancestry had been genotyped with SNP arrays and characterized for TG and HDL levels in millimoles per liter (mmol/L). The UK Biobank study was approved by the Research Ethics Committee, and informed consent was obtained from all participants. Analysis of UK Biobank data was conducted under application numbers 51436 and 26041. Type 2 diabetes designation was based on a combination of self-report or ICD-10 codes of E11.X in hospitalization or primary care records or having HbA1C level ≥ 48 mmol/mol as detailed in previous studies^11^ ^86^. Genotypes were represented by vectors of SNPs, where each SNP was encoded as 0 to represent the homozygous major allele (reference), 1 the heterozygous major/minor allele, and 2 the homozygous minor allele, yielding a participant-by-SNP genotype matrix. We performed a Bayesian regression analysis using BOLT-LMM^87^ to identify SNPs that statistically associate with log_2_(TG/HDL), normalized LDL level, or T2D status, while including sex, age, and the top 10 principal components as covariates. Significantly associated SNPs for each trait were then retained based on various p-value thresholds from 10^−4^ to 10^−8^ (**Fig. 2**, from 10^−5^ for TG/HDL due to computational limitations)

### SNP-to-Gene Mapping

Significantly associated SNPs were mapped to (potentially multiple) protein-coding genes using the union of (1) associations from cS2G^49^, (2) eQTL associations from GTEx v7^48^, or (3) the nearest gene in genomic coordinates (Genome Reference Consortium hg19 assembly^88^). For the eQTL SNP-to-gene mappings, we used adipose subcutaneous, adipose visceral omentum, liver, pancreas, adrenal gland, muscle skeletal, and uterus tissue types^42^.

### Gene Ontology Knowledge Hierarchy

The knowledge graph used for G2PT was based on the Gene Ontology^32^ Biological Process database (version 2023-07-27). This GO version was pruned using the DDOT package^89^ to include only those terms (systems) relevant to the genes with significant SNP mapping (see ‘SNP-to-Gene Mapping’ above). In particular, systems with <5 mapped genes were excluded, after which we further excluded all systems for which the annotated genes exactly matched to those of a subsystem. The final directed acyclic graph contained three types of nodes, representing SNPs, genes and systems, and three types of directed edges, representing SNP→gene mappings, gene→system annotations, and subsystem→supersystem (is_a and part_of) relations from GO.

### G2PT Model

G2PT uses a hierarchical graph transformer to integrate and distribute genotypic information across different levels in a knowledge hierarchy (here Gene Ontology as detailed above). The model uses self-attention for residual message-passing between SNPs and other biological entities in the hierarchy (systems or genes), thus propagating the effects of genetic variations on higher order biological states and translating these altered states to predict phenotypes (**Fig. 1**). Specific details are divided into the following subsections: Model Initialization and Graph Construction, Hierarchical Graph Transformer, Genetic Factor Propagation Phase, Genetic Factor Translation Phase, Model Training and Comparative Evaluation, Model Interpretation by Scoring Importance of Genes and Systems.

### Model Initialization and Graph Construction

All node embeddings—corresponding to SNPs, genes, and ontology-defined systems—were initialized randomly using uniform Xavier initialization, consistent with the model’s hidden dimension size. The attention mechanism utilizes the pre-defined connectivity between successive ontology levels to guide attention flow. During training, these hierarchical masks constrain message passing and self-attention to occur only along valid biological connections, allowing variant-level information to propagate upward through genes and systems while enabling feedback across connected subsystems.

### Hierarchical Graph Transformer

The hierarchical graph transformer (HiGT) is a modified version of a Graph Attention Transformer^29,90^ that leverages a hierarchical knowledge graph encoded into three sparse adjacency matrices (see above section ‘Gene Ontology Knowledge Hierarchy’ and ‘Model Initialization and Graph Construction’). For each node *i* with embedding state *E_i_*, the HiGT function transforms this embedding by incorporating effects from graph neighbors *j* (Eqn. 1). This transformation is computed from the neighbor embeddings via multiple attention heads Attn*_h_*, which are provided as input to a Feed Forward Neural Network (FFNN) with Layer Normalization (LN). Each head is used to perform a linear projection of the neighbor embeddings (Eqn. 2) scaled by an attention weight *α* (Eqn. 3). In particular:

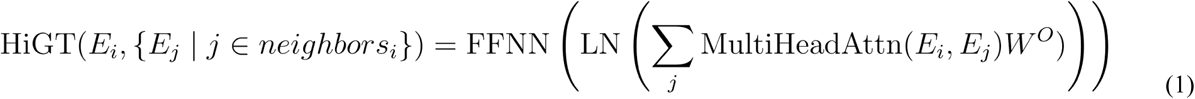

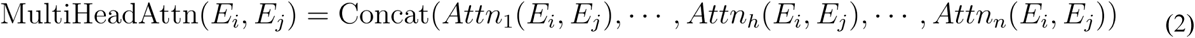

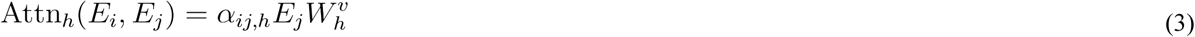

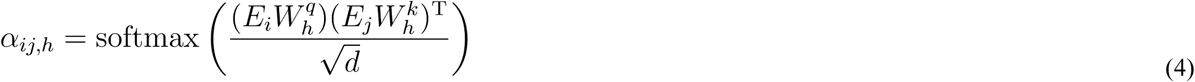

where *W^q^*, *W^k^*, *W^v^*are learnable weight matrices of dimension (*d/h)*⨉*d* encoding the central concepts of query, key, and value used in computing self-attention^29^. *W^O^* is a learnable weight matrix of dimension *d*⨉*d.* The dimension *d* encoding the size of each node embedding was set to *d*=64, the largest dimensionality that was practical given the available compute infrastructure (4 A30 GPUs). To balance complexity in a multi-headed attention model, the number of heads was set to *n*=4. The feed-forward neural networks follow the original Transformer architecture—a two-layer position-wise network with a hidden dimension four times the node embedding size (4x64) and a GeLU activation function. This HiGT function was used as the central mechanism to update the embeddings for all nodes in sequence, as described in ‘Genetic Factor Propagation’ below.

### Genetic Factor Propagation Phase

G2PT models the effects of genetic alterations via the sequential update of node states by forward propagation from SNPs to genes to systems, followed by reverse propagation from systems to genes, in a single pass. All of these updates use the HiGT formulation (eqn. 1) selecting from specific types of (target→source) edges in the knowledge graph. First, the embedding of each gene is updated given its incoming SNPs:

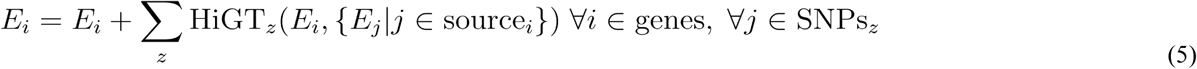

Different HiGT modules are applied based on the zygosity of the SNP alteration, z ∈ {hetero, homo_minor_}. Homozygous major SNPs represent the reference state and thus are not considered a genetic alteration. Second, the updated genes propagate their state changes upward to the systems containing these genes:

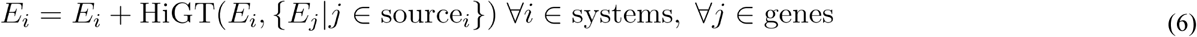

Third, state changes of systems *j* are propagated to their supersystems *i*:

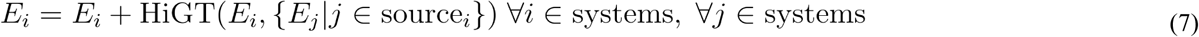

Following forward propagation, reverse propagation steps distribute the state changes downward to subsystems and genes. These reverse steps model how the particular state of a molecular system can either protect against, or expose vulnerabilities to, alterations in its component subsystems or genes. Here, state changes of systems *j* are reverse propagated to subsystems *i*:

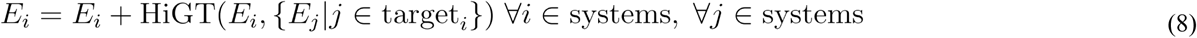

after which state changes of systems *j* are reverse propagated to genes *i*:

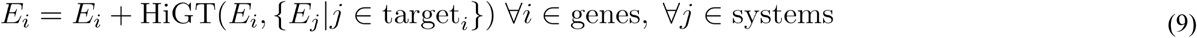

In summary, the effect of the above forward and reverse procedure is to:

(1) Forward: update all genes based on component SNPs
(2) Forward: update all systems based on component genes
(3) Forward: update all systems based on component subsystems
(4) Reverse: update all systems based on containing supersystems
(5) Reverse: update all genes based on containing systems

Overall, the HiGT functions used in each of these steps form distinct layers of the model with separately learned weights.

### Genetic Factor Translation Phase

For each participant, G2PT projects a feature vector of covariates *P^cov^* = (sex, age, PC1, …, PC10) onto an embedding of size *d* using a Multi-Layer Perceptron, yielding a participant embedding *P^embed^*.

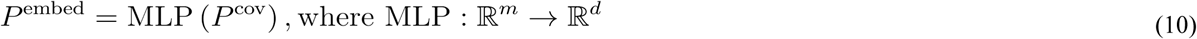

where *m* = 12 is the number of covariates. G2PT then uses the final embedding states of genes and systems (see above section ‘Genetic Factor Propagation’) to update *P^embed^*. This update occurs via the HiGT function (eqn. 1) by representing the participant as node *i* with embedding state *E_i_ = P^embed^*, and all genes and systems as graph neighbors *j* with states *E_j_*. Due to the large number of genes and systems updating a single *P^embed^*, the attention values in these operations, *α_ij,h_*, are computed using Differential Attention^91^ with a single attention head *n* = 1 to facilitate interpretation (see below section ‘Model Interpretation’). The updated embeddings from these modules are concatenated and projected through a final layer to yield *Y*, a prediction of phenotype *Y*, the log_2_(TG/HDL) phenotype. Altogether, this layer can be formulated as:

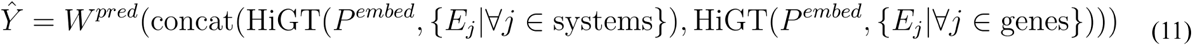

where *W^pred^*represents the learned prediction weights.

### Model Training and Comparative Evaluation

Separate models were trained for the three traits under study (TG/HDL, LDL, T2D), using nested cross-validation to train and robustly evaluate model performance in genotype-phenotype prediction. In each of five cross-validation folds, data were split into training, validation, and test sets in 3:1:1 proportion. The training set was used for selecting significant SNPs and fitting model parameters, the validation set was used for tuning model hyperparameters, and the test set comprised held-out samples for independent evaluation of performance. The model was trained to minimize the mean squared error between the true and predicted trait values, optimized with decoupled weight decay regularization (AdamW and L2 regularization)^92^. Hyperparameters were optimized through a grid search with p-value threshold set to *p* = 10^−8^ (**Supplementary Fig. 8**). Starting with the optimal baseline configuration, we further tuned the G2PT p-value threshold and training epochs, and we modified the architecture by reducing the hidden dimension and increasing the number of attention heads to manage computational costs. We also optimized parameters for other models: for XGBoost, the number of estimators, maximum depth, subsampling rate, and learning rate; and for ElasticNet, the alpha (overall strength of regularization), L1 ratio, and tolerance. For LDPred2 and Lassosum models, effect sizes of SNPs were adjusted by LD information from the training population.

### Model Interpretation by Scoring Importance of Genes and Systems

During the genetic factor translation phase (see above), G2PT assigns two sets of attention values, {*α_ik_*} and {*α_jk_*}, to weight the effects of *g_i_*∈ {*genes*} and *s_j_* ∈ {*systems*} towards the phenotype predictions of *p_k_*∈ {*participants*}. To assess the importance of these genes and systems across the population, we used the fully trained G2PT model and calculated Pearson correlations between the individual attention values and the predicted phenotype values , separately for males and females. We then averaged the absolute values of these correlations, which served as the importance score assigned to each system and gene.

### Detection of Epistatic Interactions in Important Systems

We implemented a search for epistatic interactions (**Fig. 4a**) for each system falling in the top 20 by importance (see ‘Model Interpretation by Scoring Importance’ above). For each of these systems, we first selected all SNPs mapping to genes (see ‘SNP-to-Gene Mapping’ above) with GO annotations to that system or its children. A chi-square test was used to select SNPs that exhibited significant differences in frequency between individuals with high versus low attention to the system in question (top or bottom 10% in the whole population of individuals ranked by system attention). Retained SNPs passing multiple-testing control within the system advanced to pairwise testing. SNP pairs (*SNP1*, *SNP2*) were tested for a statistical interaction in prediction of the phenotype according to a combinatorial linear model^93,94^:

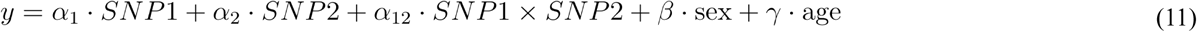

where *y* is the continuous phenotype value (TG/HDL or LDL). Each *α* represents the effect size of the corresponding SNP or SNP interaction, and *β* and *γ* represent the effect sizes of the sex and age covariates, respectively, with all *α*, *β* and *γ* values estimated from the population data. The significance of the interaction term (*α*_12_ ≄ 0) was evaluated using Wald test corrected through the Benjamini-Hochberg procedure with adjusted p-value threshold of 0.05. SNP pairs located within 1 Mb of each other were excluded in the above search, to avoid the confounding effects of linkage disequilibrium. For T2D, a binary trait, eqn. (11) above implemented logistic rather than linear regression.

### Epistasis Simulation

To evaluate G2PT’s ability to recover non-additive interactions, we simulated genotype–phenotype datasets following a sequential three-step process. First, we used a previous method^95,96^ to generate a random genotype matrix Z of dimensions *p* × *n* containing allele states across *p* individuals and *n* SNPs. We randomly designated a subset of these SNPs, *A*, to carry additive effects, and a distinct subset of SNPs to carry epistatic interactions via SNP pairs *E* (set sizes |*A*| and |*E*| were determined by hyperparameters). A vector of phenotypes *y* over all individuals was then generated by an additive-epistatic model:

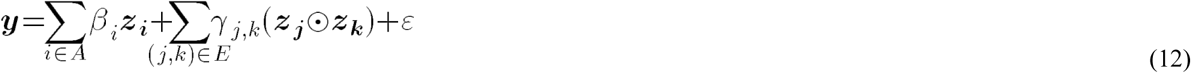

where the vector ***z_i_*** holds values of SNP *i* drawn from ***Z*** across all individuals, the scalar *β_i_* is the SNP *i* additive effect size, the scalar *γ_j,k_* is the interaction effect size between SNPs *j* and *k*, ⊙ denotes the element-wise product, and ε represents standard normal error. Effect sizes for each additive SNP and epistatic SNP pair were determined using the equations:

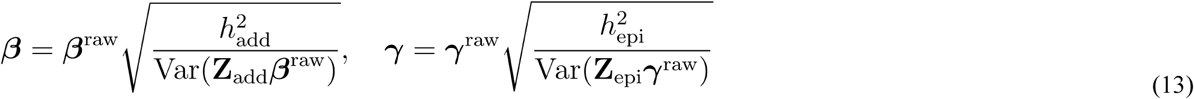

where **Z_add_**is the *p*×|*A*| sub-matrix of **Z** restricted to the additive SNP set *A*, **Z_epi_** is the *p*×|E| matrix of interaction terms formed by the element-wise products (***z_j_***⊙***z_k_***) for all epistatic pairs in *E*, and base effect size vectors (***β^raw^***, ***γ^raw^***) were drawn from a standard normal distribution and scaled by heritability hyperparameters 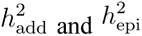 to yield the final effect size vectors (***β***, ***γ***)^97,98^.

Second, a systems ontology was constructed in a hierarchical bottom-up manner by recursively aggregating a set of initial leaf systems *S* to yield a balanced binary tree, in which the number of systems decreased by a factor of two at each successive level until a single aggregate system remained. A universe of genes *G* was created and mapped to leaf system chosen uniformly at random (set sizes |*S*| and |*G*| were determined by hyperparameters).

Third, SNPs were assigned to the ontology as follows. SNPs in the additive SNP set *A* were assigned to randomly chosen genes. SNP pairs in the epistasis set *E* were mapped 1:1 to pairs of leaf systems (*u,v*) selected as follows:

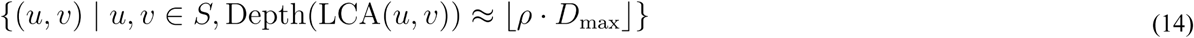

where LCA(*u*, *v*) denotes the Lowest Common Ancestor of the leaf systems *u* and *v*, and *D_max_* represents the maximum depth of the ontology (distance from the root to the leaves). The “ontology coherence” hyperparameter, *ρ*, ranges from 0 to 1, shifting (*u*,*v*) from distant systems (connected only by the ontology root) to identical leaf systems (*u* = *v*).

## Data Availability

Individual-level genomic and phenotypic data from the UK Biobank are available to researchers through an application process at https://ukbiobank.ac.uk. For this study, functional genomic annotations used for SNP-to-gene mapping were obtained in November 2023 from https://alkesgroup.broadinstitute.org/cS2G. Additionally, eQTL data were sourced from GTEx v7, accessible at https://www.gtexportal.org/home/downloads/adult-gtex/qtl. GO data were downloaded from https://release.geneontology.org/2023-07-27/index.html.

## Code Availability

The source code is available at https://github.com/idekerlab/G2PT.

## Acknowledgements

We are grateful to the National Institutes of Health for funding for this project through the following awards: Bridge2AI Common Fund (T.I.; OD032742); National Resource for Network Biology (T.I.; GM103504); National Institute of Diabetes and Digestive and Kidney Diseases (A.R.M.; DK123422) and National Heart Lung and Blood Institute (A.R.M.; HL159760). This research was also supported by the US Department of Veterans Affairs (VA, A.R.M; I01BX006293).

