## Supplementary Information for "A genotype-phenotype transformer to assess and explain polygenic risk"

### Supplementary Figures

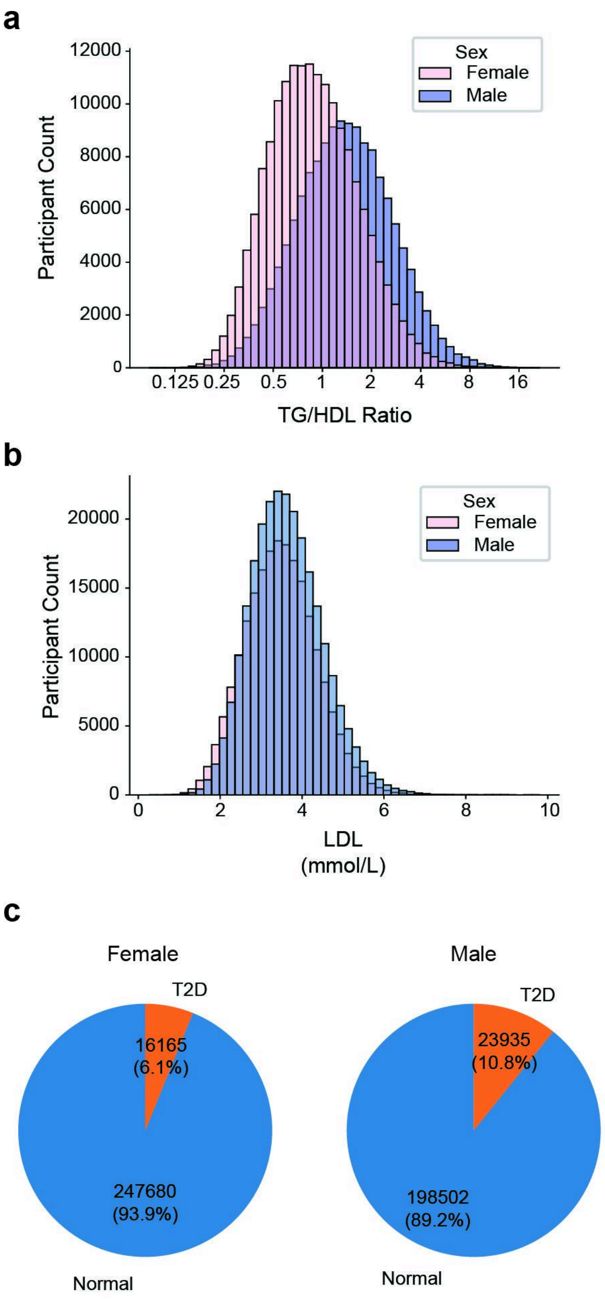


**Supplementary Fig. 1: Distribution of phenotypic trait values across individuals**. **a**, Distribution of TG/HDL ratio in the UK Biobank for female (pink) or male (blue) participants. **b**, Similar to a, for distribution of raw LDL-c (mmol/L). **c**, Pie charts of T2D (red) and normal (blue) in UK Biobank for female (left) or male (right)

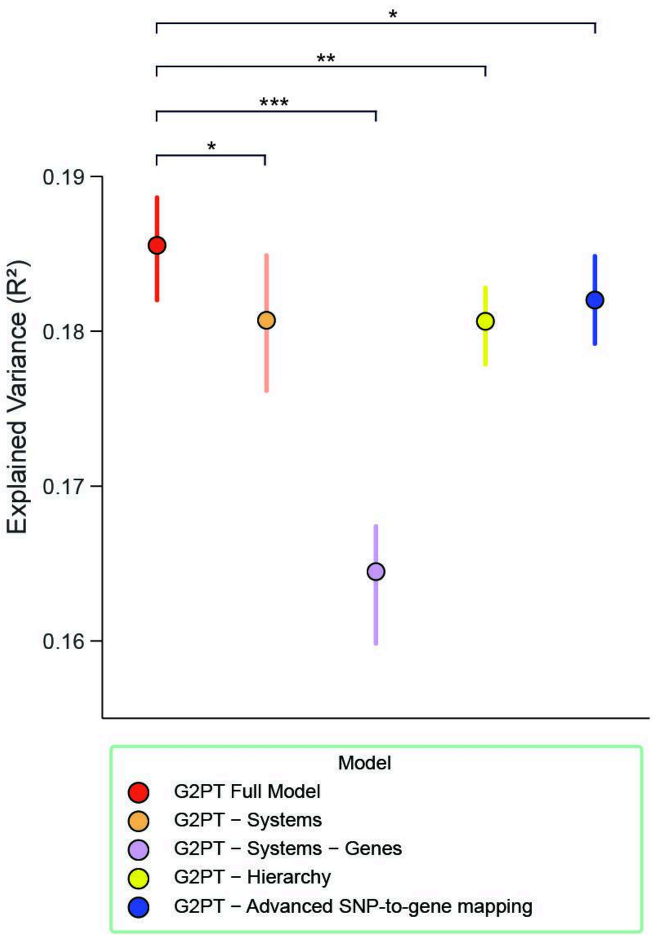


**Supplementary Fig. 2: Studies of ablated G2PT architectures.** The y-axis shows the explained variance ($R^2$) for a panel of alternative models, including the full G2PT model (red) and the model without system information (orange), without system and gene information (purple), without hierarchical connections (yellow), and without cS2G and GTEx SNP-to-gene mappings (blue). Error bars show 95% confidence intervals over five folds of cross-validation. *, significant difference in mean $R^2$ with $p < 0.05$, ** $p < 0.01$, *** $p < 0.001$.

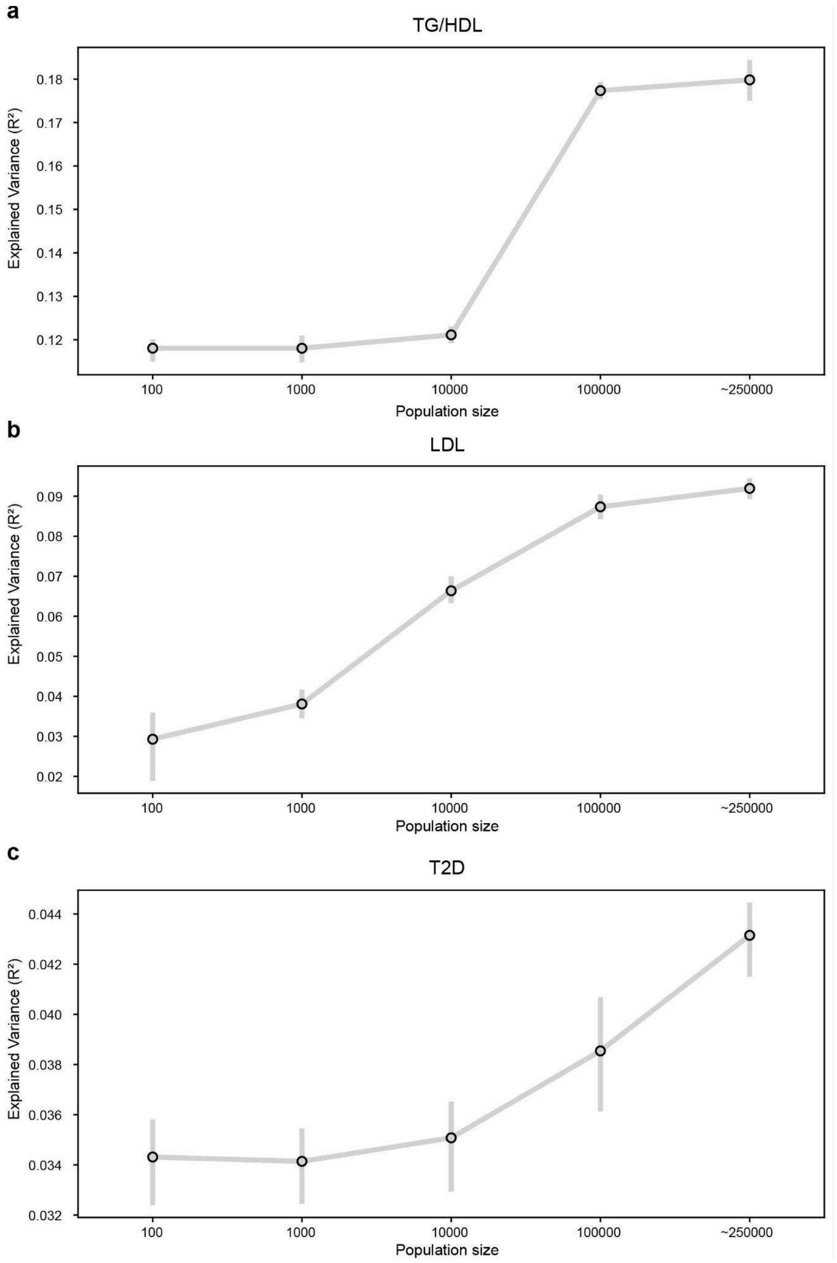


**Supplementary Fig. 3**: **Sample-size scaling of G2PT across phenotypes. a**, TG/HDL, **b**, LDL, **c**, T2D. All panels show explained variance ($R^2$) on the y-axis versus training population size on the x-axis (100, 1k, 10k, 100k, ~250k individuals). Note that ~250k represents the whole training data. Each point is the mean across nested cross-validation; error bars denote standard deviation from nested-cross validation results.

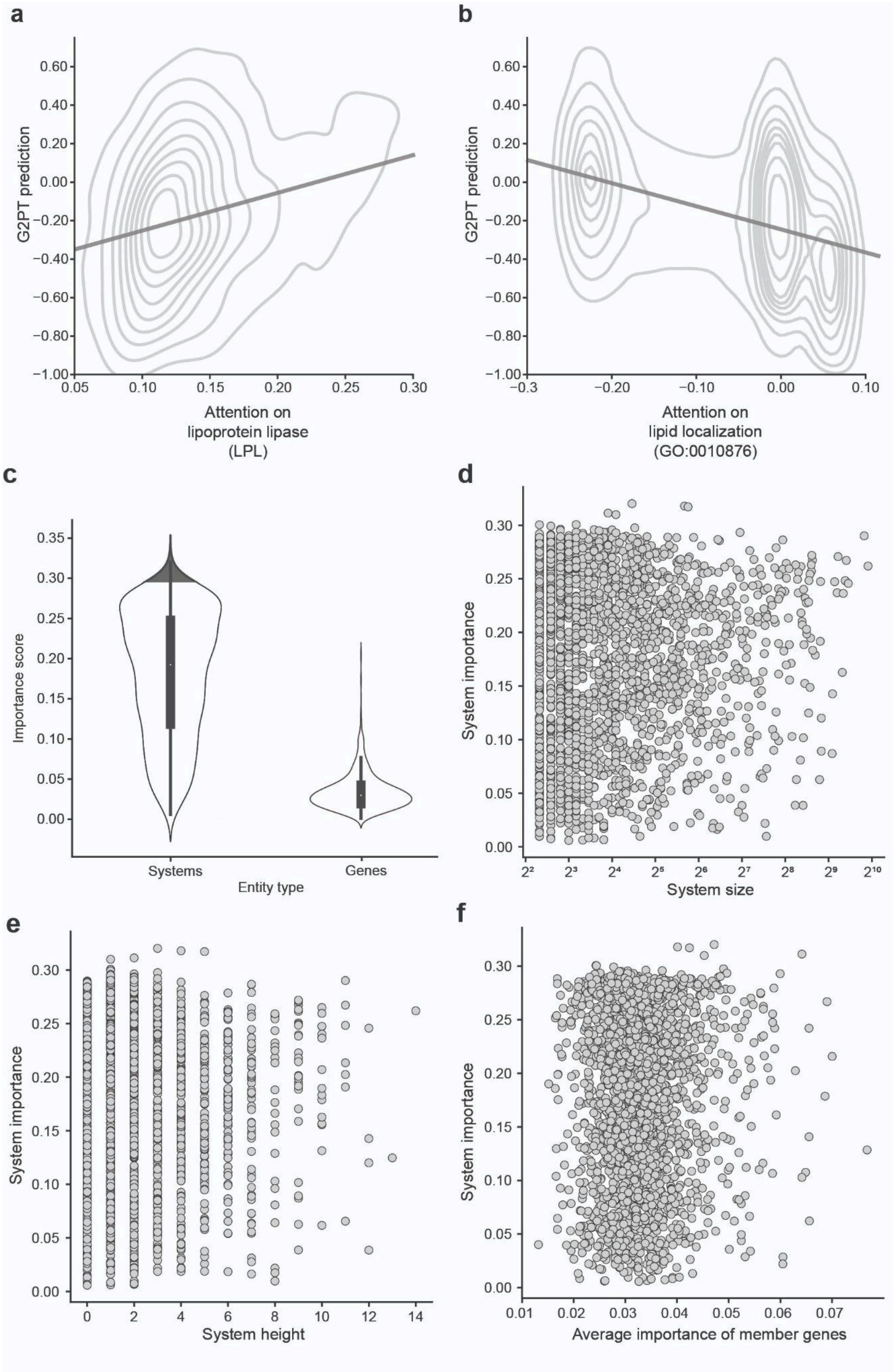


**Supplementary Fig. 4: Relationship between attention and G2PT prediction. a**, Predicted $\log_2$(TG/HDL) is plotted against Transformer attention on the lipoprotein lipase (LPL) gene over all UK Biobank individuals. The joint distribution is summarized by contours, stepping from 10% to 90% in increments of 10%. Best-fit regression line is shown. Pearson correlation with predicted TG/HDL, $r = 0.25$. **b**, Similar analysis of model attention for the sterol lipid localization system (GO:0010876). Pearson correlation with predicted TG/HDL, $r = –0.45$. **c**, Violin plots showing the distribution of importance scores for biological systems (left) and their constituent genes (right). The top shaded portion represents the 25 most important systems. The internal boxplots span the 25% to 75% quartiles, with whiskers indicating the most extreme values within 1.5 times of the interquartile range. **d-f**, Scatterplots showing the dependence of system importance score on three assorted system attributes: **d**, system size (number of genes); **e**, system height (distance in the system hierarchy from the nearest terminal leaf system); **f**, importance of member genes. Each point represents a system.

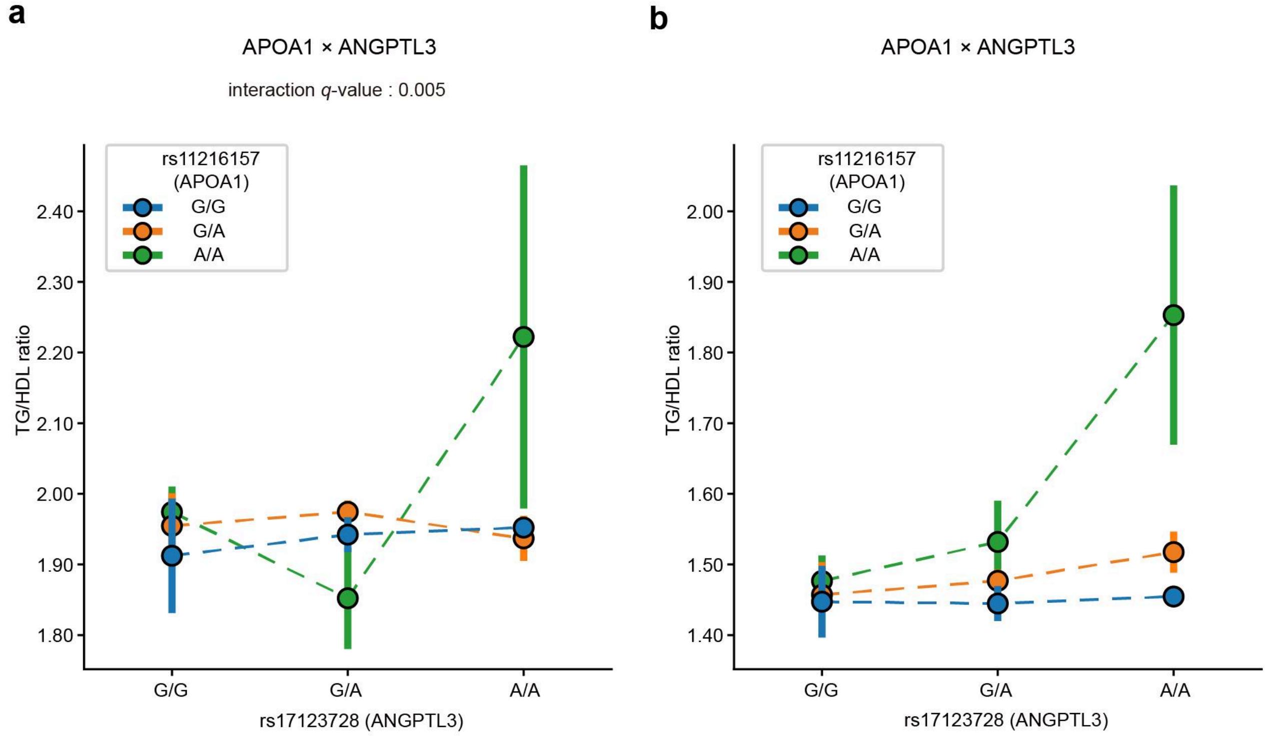


**Supplementary Fig. 5**: **Epistatic interaction reproduced in All of Us caucasian cohort**. **a**, Epistatic interaction of rs17123728 (ANGPTL3) with rs11216157 (APOA1) on measured TG/HDL ratio. Points and error bars show rs11216157 with standard error of TG/HDL ratio for subsets of individuals stratified by genotype. Colors denote rs7499892 genotype: G/G (blue, homozygous major allele), G/A (orange, heterozygous), and A/A (green, homozygous minor allele). X-axis denotes rs17123728 genotype: G/G (homozygous major allele), G/A (heterozygous), A/A (homozygous minor allele). **b**, Similar to panel (**a**), but showing the TG/HDL values predicted by the G2PT model.

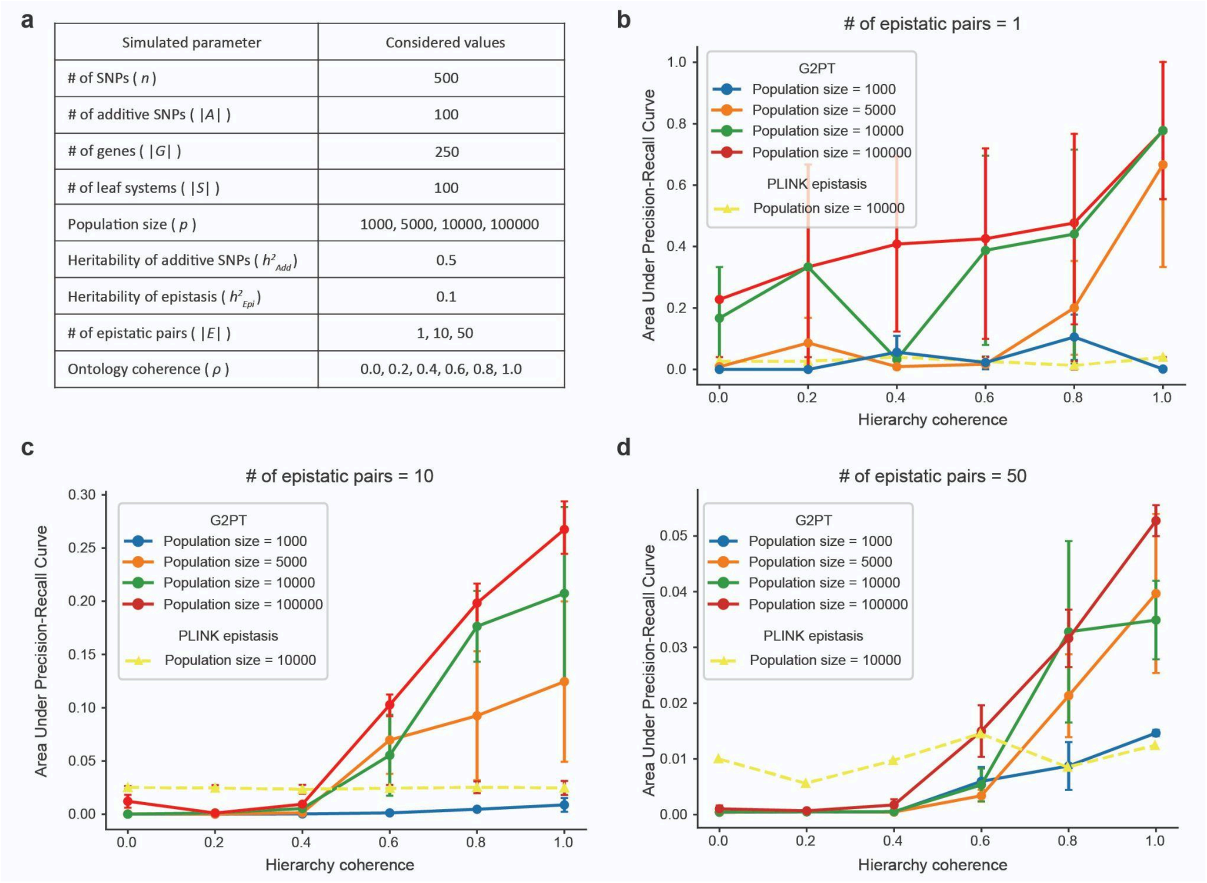

| Simulated parameter | Considered values |
|---|---|
| # of SNPs ( $n$ ) | 500 |
| # of additive SNPs ( $\|A\|$ ) | 100 |
| # of genes ( $\|G\|$ ) | 250 |
| # of leaf systems ( $\|S\|$ ) | 100 |
| Population size ( $p$ ) | 1000, 5000, 10000, 100000 |
| Heritability of additive SNPs ( $h^2_{Add}$) | 0.5 |
| Heritability of epistasis ( $h^2_{Epi}$) | 0.1 |
| # of epistatic pairs ( $\|E\|$ ) | 1, 10, 50 |
| Ontology coherence ( $\rho$ ) | 0.0, 0.2, 0.4, 0.6, 0.8, 1.0 |



**Supplementary Fig. 6**: **Supplemental epistasis simulations. a**, Table of parameters for epistatic simulation. **b**, Area under the precision–recall curve (AUPR) as a function of ontology coherence ρ and population size (1k, 5k, 10k, 100k). Circle points and error bars denote mean and standard deviation of performance across replicates. Triangle points denote performance of PLINK epistasis. **c**,**d**, Similar to panel **b** with alternative epistasis parameter settings.

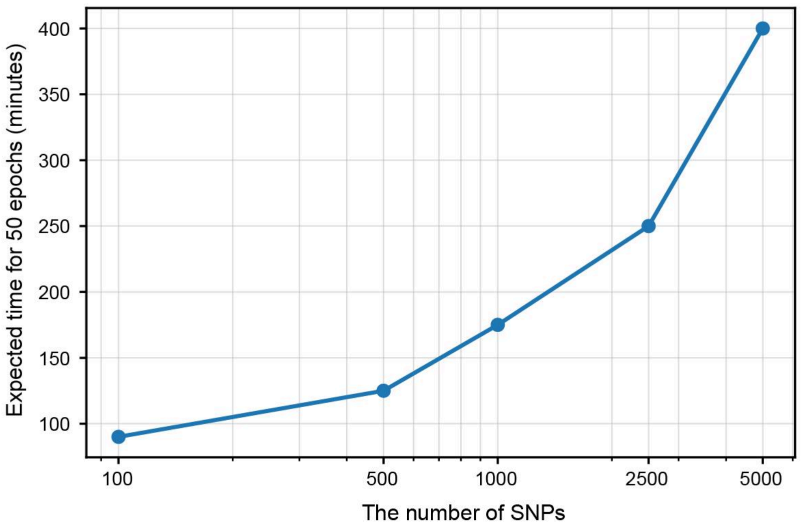


**Supplementary Figure 7: Estimated training time as a function of SNP set size.** The curve shows wall-clock minutes required to complete 50 epochs of training as the number of input SNPs increases. Training time rises exponentially.

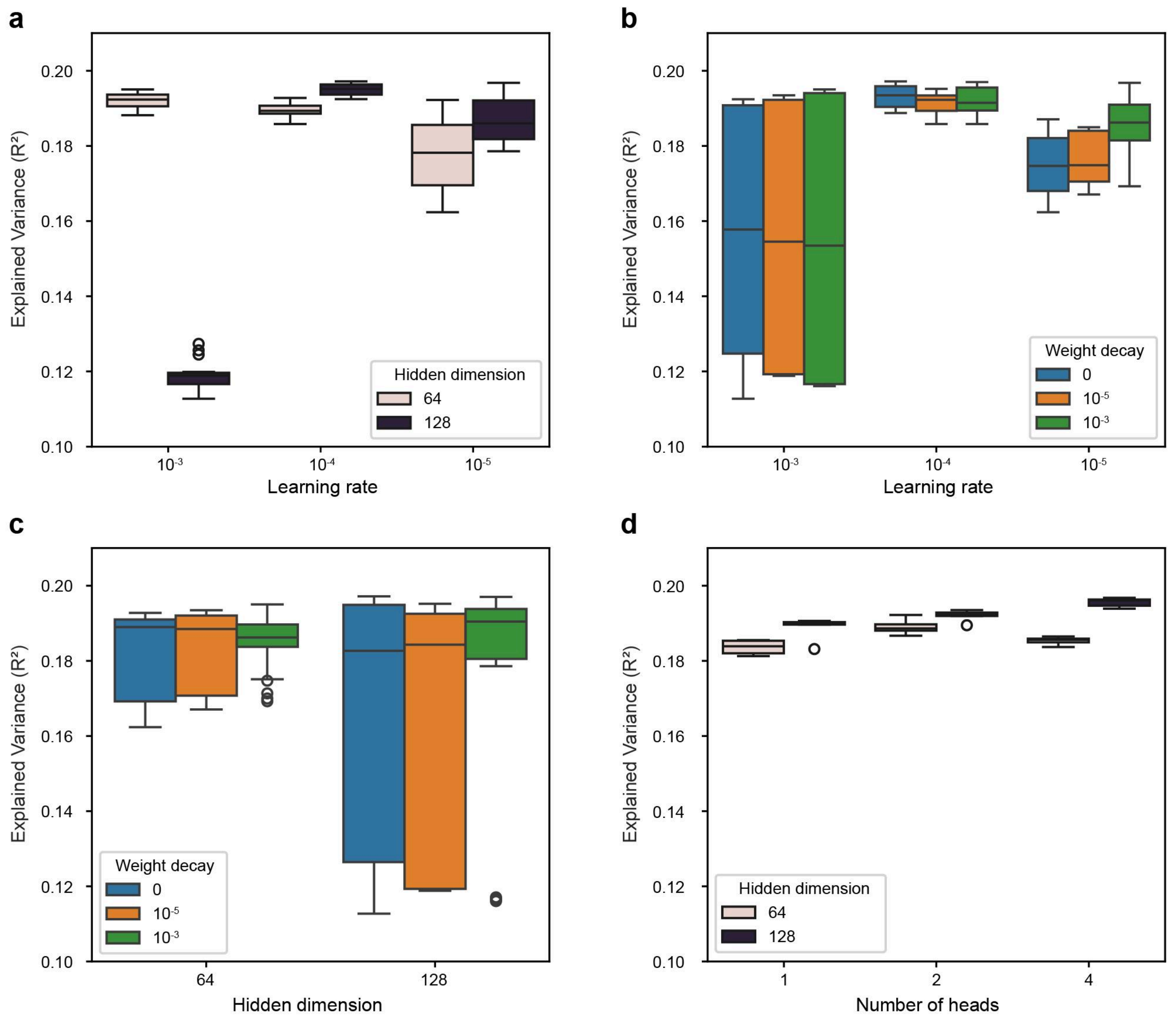


**Supplementary Figure 8: Hyperparameter settings and G2PT performance.** All panels show explained variance ($R^2$) across nested cross-validation folds. **a**, Learning rate ($10^{-3}$, $10^{-4}$, $10^{-5}$) stratified by hidden dimension (64, 128). **b**, Learning rate stratified by weight decay (0, $10^{-5}$, $10^{-3}$). **c**, Hidden dimension stratified by weight decay (0, $10^{-5}$,$10^{-3}$). **d**, Number of attention heads (1, 2, 4) stratified by hidden dimension (64, 128). Central lines show medians; boxes, interquartile ranges; whiskers indicate the most extreme values within 1.5 times of the interquartile range.

### Supplementary Tables

**Supplementary Table 1**. Number of SNPs, genes, and systems by the phenotype and p-value threshold.

| Phenotype | p-value threshold | # of SNPs | # of genes | # of systems |
|---|---|---|---|---|
| TG/HDL ratio | $10^{-5}$ | 4334 | 1176 | 1553 |
| | $10^{-6}$ | 2753 | 733 | 1091 |
| | $10^{-7}$ | 1955 | 514 | 816 |
| | $10^{-8}$ | 1501 | 392 | 653 |
| LDL | $10^{-4}$ | 1213 | 314 | 1018 |
| | $10^{-5}$ | 764 | 180 | 641 |
| | $10^{-6}$ | 537 | 123 | 477 |
| | $10^{-7}$ | 420 | 96 | 364 |
| | $10^{-8}$ | 340 | 82 | 326 |
| T2D | $10^{-4}$ | 706 | 191 | 690 |
| | $10^{-5}$ | 337 | 81 | 347 |
| | $10^{-6}$ | 206 | 47 | 225 |
| | $10^{-7}$ | 143 | 33 | 166 |
| | $10^{-8}$ | 105 | 23 | 111 |

**Supplementary Table 2.** MAGMA top-25 systems and importance scores

| System name | MAGMA importance† | G2PT importance |
|---|---|---|
| Very-low-density lipoprotein particle remodeling (GO:0034372) | 20.00 | 0.19 |
| High-density lipoprotein particle remodeling (GO:0034375) | 18.70 | 0.16 |
| Reverse cholesterol transport (GO:0043691) | 17.95 | 0.18 |
| Plasma lipoprotein particle remodeling (GO:0034369) | 13.61 | 0.15 |
| Protein-lipid complex remodeling (GO:0034368) | 13.61 | 0.33 |
| Phospholipid efflux (GO:0033700) | 13.56 | 0.28 |
| Regulation of Cdc42 protein signal transduction (GO:0032489) | 13.46 | 0.17 |
| Acylglycerol homeostasis (GO:0055090) | 11.45 | 0.25 |
| Triglyceride homeostasis (GO:0070328) | 11.45 | 0.29 |
| Cholesterol homeostasis (GO:0042632) | 10.04 | 0.31 |
| Sterol homeostasis (GO:0055092) | 10.04 | 0.26 |
| Cholesterol efflux (GO:0033344) | 9.58 | 0.26 |
| Plasma lipoprotein particle organization (GO:0071827) | 9.16 | 0.34 |
| High-density lipoprotein particle clearance (GO:0034384) | 9.11 | 0.06 |
| Regulation of lipoprotein lipase activity (GO:0051004) | 9.04 | 0.19 |
| Regulation of plasma lipoprotein particle levels (GO:0097006) | 9.01 | 0.23 |
| Triglyceride catabolic process (GO:0019433) | 8.56 | 0.27 |
| Regulation of cholesterol storage (GO:0010885) | 8.53 | 0.31 |
| Positive regulation of cholesterol transport (GO:0032376) | 8.45 | 0.13 |
| Phospholipid homeostasis (GO:0055091) | 8.20 | 0.29 |
| † MAGMA importance score is from p-value of system significance |  |  |

**Supplementary Table 3**. Significant epistatic interactions identified in top 20 important systems.

| System name | SNP 1 | Gene 1† | SNP 2 | Gene 2† | Adjusted p-value‡ |
|---|---|---|---|---|---|
| Phospholipid transport (GO:0015914) | rs11216169 | APOA4 | rs7499892 | CETP | 0.048992 |
| | rs11216169* | APOA4 | rs1800775* | CETP | 0.046611 |
| | rs11216169 | APOA4 | rs7205804 | CETP | 0.048992 |
| | rs11216169 | APOA4 | rs1532624 | CETP | 0.048992 |
| | rs11216169 | APOA4 | rs1864163 | CETP | 0.048992 |
| | rs3764261 | CETP | rs4420638 | APOC1 | 0.048992 |
| | rs247617 | CETP | rs4420638 | APOC1 | 0.048992 |
| | rs11216169 | APOA4 | rs1800775 | CETP | 0.039122 |
| Cholesterol transport (GO:0030301) | rs11216169 | APOA4 | rs13089415 | PPARG | 0.039122 |
| | rs11216169* | APOA4 | rs1800775* | CETP | 0.049193 |
| Protein dephosphorylation (GO:0006470) | rs4802113 | TGFB1 | rs910619 | FKBP1A | 0.033128 |
| Lipid storage (GO:0019915) | rs11216157 | APOA1 | rs17123728 | ANGPTL3 | 0.000777 |
| Protein-lipid complex remodeling (GO:0034368) | rs2083637 | LPL | rs1145187 | APOA5 | 0.039342 |
| | rs2083637 | LPL | rs1145189 | APOA5 | 0.039342 |
| | rs3916027 | LPL | rs1145189 | APOA5 | 0.039342 |
| | rs12682115 | LPL | rs4420638 | APOC1 | 0.039342 |
| | rs4922117 | LPL | rs1145187 | APOA5 | 0.039342 |
| | rs4922117 | LPL | rs1145189 | APOA5 | 0.039342 |
| | rs3735964 | LPL | rs4420638 | APOC1 | 0.039342 |
| | rs1059611 | LPL | rs4420638 | APOC1 | 0.039342 |
| | rs17489282 | LPL | rs1145187 | APOA5 | 0.039342 |
| | rs17489282 | LPL | rs1145189 | APOA5 | 0.039342 |
| | rs1919484 | LPL | rs1145187 | APOA5 | 0.039342 |
| | rs1919484 | LPL | rs1145189 | APOA5 | 0.039342 |
| | rs17411126 | LPL | rs1145187 | APOA5 | 0.039342 |
| | rs17411126 | LPL | rs1145189 | APOA5 | 0.039342 |
| | rs17482753 | LPL | rs4420638 | APOC1 | 0.039342 |
| | rs331 | LPL | rs1145187 | APOA5 | 0.039342 |
| | rs331 | LPL | rs1145189 | APOA5 | 0.039342 |
| | rs327 | LPL | rs1145187 | APOA5 | 0.039342 |
| | rs765547 | LPL | rs1145187 | APOA5 | 0.039342 |
| | rs297 | LPL | rs1145187 | APOA5 | 0.039342 |
| | rs17411045 | LPL | rs1145187 | APOA5 | 0.039342 |
| | rs327 | LPL | rs1145189 | APOA5 | 0.039342 |
| | rs10503669 | LPL | rs4420638 | APOC1 | 0.039342 |
| | rs765547 | LPL | rs1145189 | APOA5 | 0.039342 |
| | rs297 | LPL | rs1145189 | APOA5 | 0.039342 |
| | rs12678919 | LPL | rs4420638 | APOC1 | 0.039342 |
| | rs7816447 | LPL | rs4420638 | APOC1 | 0.039342 |
| | rs17411045 | LPL | rs1145189 | APOA5 | 0.039342 |
| | rs12679834 | LPL | rs4420638 | APOC1 | 0.039342 |

† Both gene 1 and 2 are annotated members of the relevant GO system from column 1.
‡ Statistical regression model, test of significant interaction term with Benjamini-Hochberg correction. All SNP pairs with adjusted $p < 0.05$ shown.
* Epistatic interaction between rs11216169 and rs1800775 exists in both phospholipid transport and cholesterol transport system.
